# The need for ecologically realistic studies on the health effects of microplastics

**DOI:** 10.1101/2022.11.21.517421

**Authors:** C Lauren Mills, Joy Savanagouder, Marcia de Almeida Monteiro Melo Ferraz, Michael J Noonan

**Affiliations:** The Irving K. Barber Faculty of Science, The University of British Columbia, Okanagan Campus; Clinic of Ruminants, Faculty of Veterinary Medicine, Ludwig-Maximilians University of Munich, Sonnenstr. 16, Oberschleissheim, 85764, Germany

**Keywords:** Microplastics, terrestrial, health, biomimetic

## Abstract

Plastic pollution is now so widespread that microplastics are consistently detected in every biological sample surveyed for their presence. Despite their pervasiveness, very little is known about the effects of microplastics on the health of terrestrial species. While emerging studies are showing that microplastics represent a potentially serious threat to animal health, data have been limited to *in vivo* studies on laboratory rodents that were force fed plastics. The extent to which these studies are representative of the conditions that animals and humans might actually experience in the real world is largely unknown. Here, we review the peer-reviewed literature in order to understand how the concentrations and types of microplastics being administered in lab studies compare to those found in terrestrial soils. We found that lab studies have heretofore fed rodents microplastics at concentrations that were hundreds of thousands of times greater than they would be exposed to in nature. Furthermore, health effects have been studied for only 10% of the microplastic polymers that are known to occur in soils. The plastic pollution crisis is arguably one of the most pressing ecological and public health issues of our time, yet existing lab-based research on the health effects of terrestrial microplastics does not reflect the conditions that free-ranging animals are actually experiencing. Going forward, performing more true-to-life research will be of the utmost importance to understand the impacts of microplastics and maintain the public’s faith in the scientific process.

## 1. Introduction

The invention of plastics in the early 1900s revolutionized human societies (Thompson et al., 2009), yet the excessive consumption of short-lived and single-use plastics has resulted in plastics accumulating almost everywhere on Earth (Cole et al., 2011; Rochman & Hoellein, 2020). Plastic pollution is now so widespread that microplastics – plastic particles between 0.1 μm and 5 mm – are consistently detected in every biological sample surveyed for their presence (Duis & Coors, 2016; Bergami et al., 2020). The ubiquitous and long-lived nature of microplastics makes them a worrying environmental contaminant, yet, despite their pervasiveness, very little is known about how microplastics might be impacting the health of species living in terrestrial ecosystems. This stands in stark contrast to the fact that 80% of species live on land (Grosberg et al., 2012), and that the volume of microplastics in terrestrial systems may be greater than that in oceans (de Souza Machado et al., 2012; Hurley & Nizzetto, 2018).

Though evidence is still extremely limited, emerging studies are showing that microplastics represent a potentially serious threat to the health of terrestrial species, and may impact an array of biological functions (Huang et al., 2022; Lou et al., 2019). For instance, recent work in mice and rats has demonstrated the detrimental effects of microplastics on sperm production (Jin et al., 2021). Similarly, a study conducted by Wang et al. (2022) indicated that mice exposed to MPs experienced both necroptosis and inflammation within bladder epithelium, while Djouina et al. (2022) found that microplastics can adversely affect the small intestine and colon of mice, causing histological and immune disturbances, as well as inflammation. Data have been limited to *in vivo* studies on laboratory rodents that were force fed plastics, however, and there are currently no studies describing the health effects of microplastics exposure outside of laboratory settings. Thus, although the findings from these studies are certainly worrying, the extent to which they are representative of the conditions that humans and animals are actually experiencing in the real world is largely unknown. Here, we review the peer-reviewed literature to explore the extent to which lab studies on the effects of microplastics are representative of the conditions that animals are experiencing in the real world. In particular we focused on understanding how the concentrations of microplastics and types of polymers being administered in lab studies compared to those found in terrestrial soils. Although our focus was on microplastics in soils, this is not the only path of exposure to microplastics. For instance, plants can uptake microplastics (Azeem et al., 2021), which can then be ingested by herbivorous/omnivorous species. Airborne microplastics can also be inhaled, with intake rates that may be comparable to ingestion (Cox et al., 2019). Most studies on airborne microplastics quantify concentrations in terms of deposition rates (Sridharan et al., 2021), however, making direct comparisons to lab studies impossible, and there is little information on the microplastic exposure and ingestion rates of free-ranging terrestrial species. Nonetheless, air and waterborne microplastics will ultimately accumulate in soils (Guo et al., 2020, Sridharan et al., 2021), and soils are at the base of many terrestrial food webs (de Souza Machado et al., 2018). The concentrations of microplastics in soils are thus likely to be broadly representative of exposure levels. Our results can help provide much needed context to the findings of existing health studies, as well as an ecologically relevant baseline that can help guide future lab studies on the health effects of terrestrial microplastics.

## 2. Materials and methods

We first identified studies from the peer-reviewed literature that were focused on the health effects of microplastics on terrestrial animals, or on microplastics in terrestrial soil environment via a Google Scholar search for the terms “microplastics”, “microplastics” and “mice”, “microplastics” and “rats”, “microplastics” and “rodents”, “microplastics in lab”, and “microplastics in soil”. Any *in vivo* lab studies not directly relating to the ingestion of microplastics were excluded as they were beyond the scope of our effort. Similarly, studies where soil samples were taken from lakes or river beds were excluded as our focus was on describing the conditions being experienced by terrestrial species. Through this initial search, a total of 93 peer-reviewed studies were compiled; 55 studies focused on microplastics in *in vivo* lab studies, and 38 focused on microplastics in terrestrial soil environments. For *in vivo* studies we extracted information on the polymer type, concentrations fed to laboratory rodents, and diameter, volume, and density of the microplastic particles. The microplastic type and final concentrations found in the soil environment were extracted from soil studies. There was very little consistency in the units across studies, and so to standardize microplastic measurements, all concentrations were converted to items/kg. To do this, polymer type was required to identify the density of the plastic, while diameter was required to calculate the volume. The known volume, density, and concentrations were then used in conjunction to calculate the number of particles and convert the data to items/kg. If any information required to make this conversion was absent from a study, it was excluded from subsequent analyses. Similarly, soil studies were excluded if information on the concentrations of microplastic were absent, or if they were experimental studies. This further narrowed the number of studies down to a total of 28 *in vivo* studies describing 67 experimental concentrations, and 22 soil studies with data on 48 sites.

## 3. Results and discussion

The median concentration of microplastics fed to laboratory rodents in *in vivo* studies was 36,841,422 items/kg. This was over 78,000 times greater than the median concentration of 471 items/kg found in soil (Fig. 1A). The highest recorded concentration of microplastics in any soil sample was 18,760 items/kg which was found in agricultural soil along China’s Chai river valley (Zhang & Liu, 2019); only 5 out of the 28 compiled lab studies used concentrations below this amount. We also found that while 28 different plastic polymers have been found to occur in soil, the health effects of only 3 polymers have been studied to date, with the overwhelming majority of *in vivo* experiments having focused on polystyrene (Fig. 1B). The stark contrast between the types and concentrations of microplastics being administered to lab rodents in *in vivo* studies versus the conditions these animals are likely to encounter in the wild questions the utility of these findings and illustrates the need for more ecologically realistic studies.

**Figure 1.**
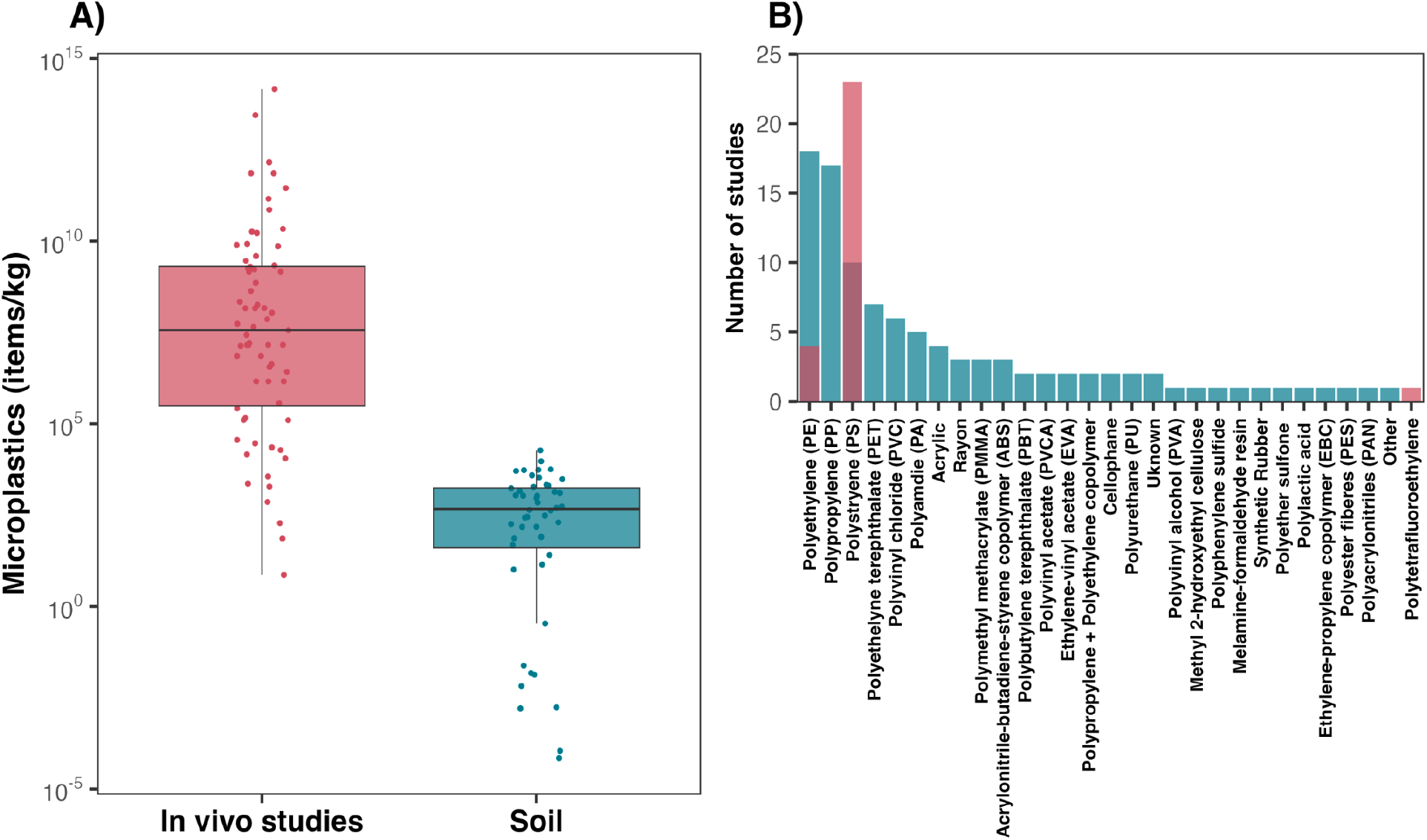
The boxplot in A shows the concentrations of MPs fed to rodents in *in vivo* lab studies, compared to those of MPs found in soils. In B the number of soil studies which identified different plastic polymers are shown in blue, whereas the number of polymers assessed via *in vivo* health studies are shown in red. Data were compiled from 50 peer-reviewed studies; 22 on MPs in soil and 28 on the health effects of MPs.

Notably, and in light of this disconnect, a common trend across lab studies was the lack of any rationale for the concentrations of microplastic that were administered. The 11 studies that did provide justification chose concentrations that were based either on the concentrations of microplastic found in rivers (Liu et al., 2022), or on existing *in vivo* studies (Choi et al., 2021, Hou et al., 2021; Li et al., 2020; Lou et al., 2019; Mu et al., 2022; Shi et al, 2022; Wang et al., 2022; Wang et al., 2022; Yang et al., 2019; Yang et al., 2022). For instance, Yang et al. (2019) and Mu et al. (2022), both based their study designs on work on mice by Deng et al. (2017). Deng et al. (2017) which, however, based their study on MP concentrations found in rivers, and therefore it does not accurately depict terrestrial environments. Thus, while a handful of lab studies did provide some form of justification for their study design, the extent to which these studies are representative of the conditions that humans and animals are actually experiencing in the real world is questionable.

## 4. Conclusions

The plastic pollution crisis is arguably one of the most pressing ecological and public health issues of our time, yet existing research on the health effects of terrestrial microplastics does not accurately reflect the conditions that humans and animals are actually experiencing. Paired with this disconnect is the fact that 1,196 animals were sacrificed to generate the findings of these 28 studies, yet the majority of these animals were fed tens to hundreds of thousands of times more plastic than they would ever be exposed to in the wild. Because microplastics research also receives frequent media attention, performing true-to-life studies is of the utmost importance so as to not erode the public’s faith in the scientific process. It therefore falls on the scientific community to describe the ecologically realistic effects of microplastics on the health of terrestrial species in order for well-founded mitigation efforts to be launched. Going forward, performing more true-to-life research will be of the utmost importance to understand the impacts of microplastics and maintain the public’s faith in the scientific process.

## Acknowledgments

This work was supported by an NSERC Discovery Grant RGPIN-2021-02758 to MJN, as well as the Canadian Foundation for Innovation. MAMMF was supported by LMUexcellent, funded by the Federal Ministry of Education and Research (BMBF) and the Free State of Bavaria under the Excellence Strategy of the Federal Government and the Länder.

## Author contributions

MJN and MAMMF conceived of the study, CLM and JS conducted the literature review, CLM and MJN wrote the first manuscript draft, and all co-authors assisted with writing the final version of the manuscript.

## Competing interests

Authors declare that they have no competing interests.

## Data and materials availability

The data and R scripts used to carry out this study are openly available on GitHub at https://github.com/QuantitativeEcologyLab/MP_Disconnect.

## Notes

https://github.com/QuantitativeEcologyLab/MP_Disconnect

### Competing Interest Statement

The authors have declared no competing interest.

